# Exact Topological Inference for Paired Brain Networks *via* Persistent Homology

**DOI:** 10.1101/140533

**Authors:** Moo K. Chung, Victoria Villalta-Gil, Hyekyoung Lee, Paul J. Rathouz, Benjamin B. Lahey, David H. Zald

## Abstract

We present a novel framework for characterizing paired brain networks using techniques in hyper-networks, sparse learning and persistent homology. The framework is general enough for dealing with any type of paired images such as twins, multimodal and longitudinal images. The exact nonparametric statistical inference procedure is derived on testing monotonic graph theory features that do not rely on time consuming permutation tests. The proposed method computes the exact probability in quadratic time while the permutation tests require exponential time. As illustrations, we apply the method to simulated networks and a twin fMRI study. In case of the latter, we determine the statistical significance of the heritability index of the large-scale reward network where every voxel is a network node.

## 1 Introduction

There are many studies related to paired images: longitudinal studies with two repeat scans [7], multimodal imaging study involving PET and MRI [13] and twin imaging studies [15]. The paired images are usually analyzed by relating voxel measurements that match across two images in a mass univariate fashion at each voxel. Compared to the paired image setting, paired brain networks have not been often analyzed possibly due to the lack of problem awareness and analysis frameworks.

In this paper, we present a new unified statistical framework that can integrate paired networks in a holistic fashion by pairing every possible combination of voxels across two images. This is achieved using a hyper-network that connects multiple smaller networks into a larger network. Although hyper-networks are frequently used in machine learning [16], the concept has not been often used in medical imaging. Jie et al. used the hyper-network framework from the resting-state fMRI in classifying MCI from AD in a machine learning framework [9]. Bezerianos et al. constructed the hyper-network from the coupling of EEG activity of pilots and copilots operating an aircraft albeit using ad-hoc procedures without an explicit model specification [1]. Motivated particularly by the science of [1], we rigorously formulate the problem of characterizing paired networks as a single hyper-network with physically nonexistent hyper-edges connecting between two existing networks.

Graph theory is an often used framework for analyzing brain networks mainly due to easier accessibility and interpretation. Since we do not know the exact statistical distribution of many graph theory features, resampling techniques such as the permutation tests have been mainly used in estimating the null distribution and computing *p*-values. The availability of inexpensive and fast computers made the permutation tests a natural choice for computing *p*-values when underlying distributions are unknown. However, even with fast computers, permutation test is still extremely slow even for small-scale problems. There is a strong need for fast inference procedures. We propose a new exact inference procedure for graphs and networks. The method speeds up the computation by counting the number of permutations combinatorially instead of empirically computing them by numerically generating large number of permutations.

Our main contributions are as follows.

1. The new formulation of the hyper-network approach for paired images. We show that hyper-networks can be effectively used as a baseline model for paired twin fMRI. The proposed holistic framework is applied to a twin fMRI study in determining the statistical significance of the brain network differences and the heritability of the reward network.
2. Showing that the topological structure of the sparse hyper-network has a monotonic nestedness. This is used to define monotonic graph features and subsequently in computing *topologically aware* distance between networks.
3. The derivation of the probability distribution of the combinatorial test procedure on graph theory feature vectors that do not rely on resampling techniques such as the permutation tests. While the permutation tests require exponential run time, our exact combinatorial approach requires quadratic run time.

## 2 Sparse hyper-network

Consider a collection of *n* paired images

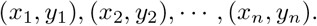

We assume *x_k_* and *y_k_* are related either genetically (twin or sibling images), scan-wise (multimodal images of the same subject) or longitudinally. We further assume (*x_k_*, *y_k_*) are independently but identically distributed for different *k*. Let **x** = (*x*_1_, ⋯, *x_n_*)^⊤^ and **y** = (*y*_1_, ⋯, *y_n_*)^⊤^ be the vectors of images. We set up a hyper-network by relating the paired vectors at different voxels *v_i_* and *v_j_*:

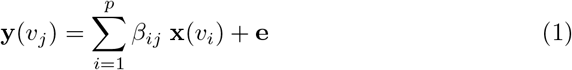

for some zero-mean noise vector **e** (Fig. 1). The parameters *β* = (*β_ij_*) are the weights of the hyper-edges between voxels *v_i_* and *v_j_* that have to be estimated. We are constructing a *physically nonexistent artificial network* across different images. For fMRI, (1) requires estimating over billions of connections, which is computationally challenging. In practice however, each application will likely to force *β* to have a specific structure that may reduce the computational burden. For this study, we will consider a reduced model relevant to our genetic imaging application:

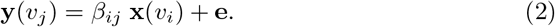

**Fig. 1:**
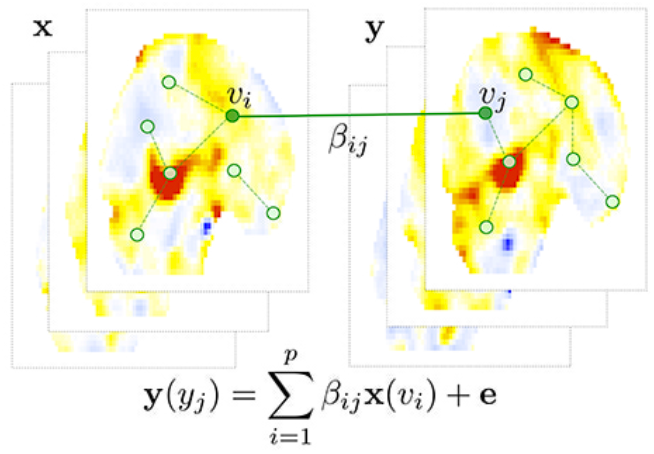
The schematic of hypernetwork construction on paired image vectors **x** and **y**. The image vectors **y** at voxel *v_j_* is modeled as a linear combination of the first image vector **x** at all other voxels. The estimated parameters *β_ij_* give the hyper-edge weights.

The scientific motivation for using the reduced model will be explained in section 6. Without loss of generality, we can center and scale **x** and **y** such that

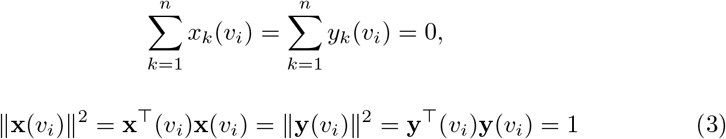

for all voxels *v_i_*.

In many applications, we have significantly larger number of voxels (*p*) than the number of images (*n*), so model (2) is an under-determined system and belongs to the *small-n large-p problem* [5]. It is necessary to regularize the model using a sparse penalty:

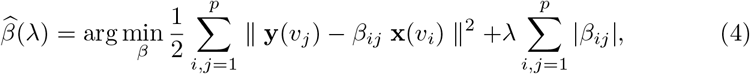

where sparse parameter λ ≥ 0 modulates the sparsity of the hyper-edges. The estimation 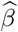 is a function of λ. When λ = 0, (4) is a least-squares problem and we obtain 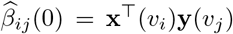, which is referred to as the *cross-correlation*. The cross-correlation is invariant under the centering and scaling.

## 3 Distance between networks

To study the topological characteristic of the hyper-network, the estimated edge weights 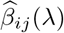 are binarized by assigning value 1 to any nonzero weight and 0 otherwise. Let *G*_λ_ be the resulting binary graph, i.e., unweighted graph, with adjacency matrix *A*_λ_ = (*a_ij_*(λ)):

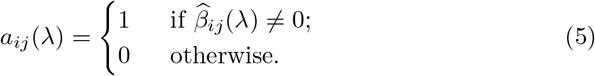

As λ increases, *G*_λ_ becomes smaller and nested in a sense that every edge weight gets smaller. This is *not* true for the binarization of other sparse network models. Our sparse hyper-network has this extra layer of additional structure that guarantees the nestedness.

### Theorem 1.

*The binary graph G*_λ_ *obtained from sparse model (4) satisfies*

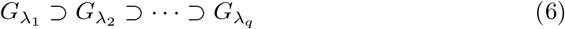

*for any* 0 ≤ λ_1_ ≤ λ_2_ ≤ ⋯ ≤ λ_*q*_ ≤ 1.

*Proof*. By solving 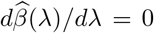 for λ ≠ 0 and considering 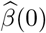 separately, it can be algebraically shown that the solution of the optimization problem (2) is given by [5]

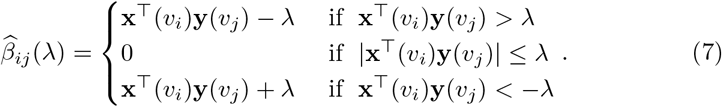

From (3) and the Cauchy-Schwarz inequality, we have |**x**^⊤^(*v_i_*)**y**(*v_j_*)| ≤ 1. Thus, if λ > 1, 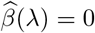. To avoid the trivial case of zero edge weights everywhere, we need 0 ≤ λ ≤ 1. From (7), we have 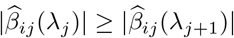 for 0 ≤ λ_*j*_ ≤ λ_*j*+1_ ≤ 1. Subsequently, from (5), *a_ij_*(λ_*j*_) ≥ *a_ij_*(λ_*j*+1_). Therefore, we have *G*_λ_*j*__ ⊃ *G*_λ_*j*+1__. □

The sequence of nested multi-scale graph structure (6) is called the *graph filtration* in persistent homology [11]. The graph filtration can be quantified using monotonic function *B* satisfying

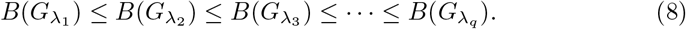

The number of connected components, which is the zero-th Betti number *β*_0_ and the most often used topological invariant in persistent homology, satisfies this condition. The size of the largest connected component satisfies a similar but opposite relation of monotonic decrease.

Given two different graph filtrations 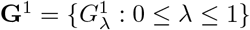 and 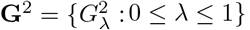, define the distance between them as

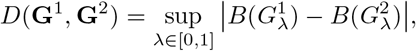

which can be discretely approximated as

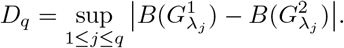

If we choose enough number of *q* such that λ_*j*_ are all the sorted edge weights, then *D*(**G**^1^, **G**^2^) = *D_q_*. This is possible since there are only up to *p*(*p* − 1)/2 number of unique edge weights in a graph with *p* nodes and 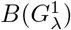 and 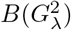 increase discretely. In practice, *ρ_j_* = *j*/*q* is chosen uniformly in [0, 1] or a divide-and-conquer strategy can be used to do adaptively grid the unit interval.

*D* satisfies all the axioms of a metric except identity. *D*(**G**^1^, **G**^2^) = 0 does not imply **G**^1^ = **G**^2^. Thus, *D* is *not* a metric. However, it will be shown in section 4 that probability *P*(*D*(**G**^1^, **G**^2^) = 0) = 0 showing such event rarely happens in practice. Thus, *D* can be treated as a metric-like in applications without much harm.

## 4 Exact topological inference

We are interested in testing the null hypothesis *H*_0_ of the equivalence of two monotonic graph feature functions:

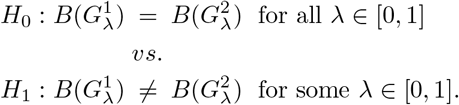

As a test statistic, we propose to use *D_q_*. The test statistic takes care of the multiple comparisons by the use of supremum. Under the null hypothesis, we can derive the probability distribution of *D_q_* combinatorially without numerically permuting images.

### Theorem 2.

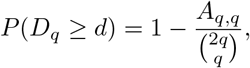

*where A_u,v_ satisfies A_u,v_* = *A*_*u*–1,*v*_ + *A*_*u,v*–1_ *with the boundary condition A*_0,*q*_ = *A*_*q*,0_ = 1 *within band* |*u* − *v*| < *d*.

*Proof*. The proof is similar to the combinatorial construction of Kolmogorov-Smirnov (KS) test [2,5,8]. Combine two monotonically increasing vectors

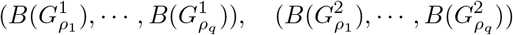

and arrange them in increasing order. Represent 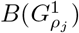 and 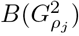 as ↑ and → respectively. For example, ⇈→↑→→ ⋯. There are exactly *q* number of ↑ and *q* number of → in the sequence. Treat the sequence as walks on a Cartesian grid. → indicates one step to the right and ↑ indicates one step up. Thus the walk starts at (0, 0) and ends at (*q, q*) (Fig. 2). There are total 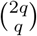 possible number of paths, which forms the sample space. This is also the total number of possible permutations between the elements of the two vectors. The null assumption is that the two vectors are identical and there is no preference to one vector element to another. Thus, each walk is equally likely to happen in the sample space.

**Fig. 2:**
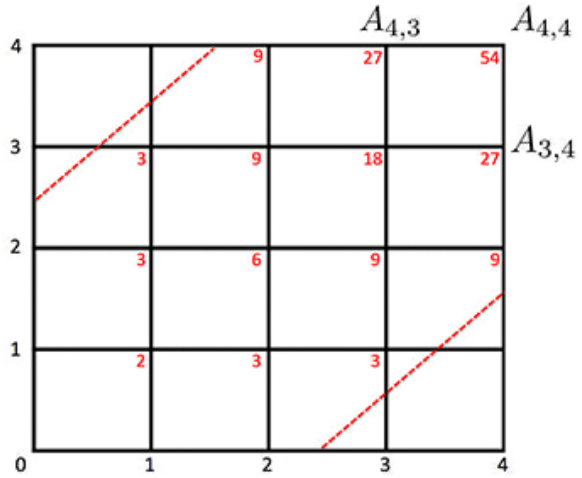
*A_u,v_* are computed within the boundary (dotted red line). The red numbers are the number of paths from (0, 0).

The values of *B*(*G*^1^_*ρ_j_*_) and *B*(*G*^2^_*ρ_j_*_) satisfying the condition sup_j_ |*B*(*G*^1^_*ρ_j_*_) − *B*(*G*^2^_*ρ_j_*_)| < *d* are the integer coordinates of (*u, v*) on the path satisfying |*u* − *v*| < *d*. These are integer grid points within the boundary *v* = *u* ± *d* (dotted red lines in Fig. 2). Thus the probability can be written as

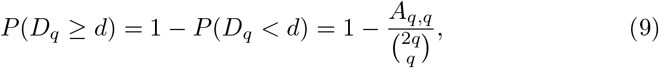

where *A_u,v_* be the total number of passible paths from (0, 0) to (*u, v*) within the boundary. Since there are only two paths (either ↑ or →), *A_u,v_* can be computed recursively as

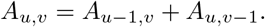

within the boundary. On the boundary, *A*_0,*q*_ = *A*_*q*,0_ = 1 since there is only one path. □

Theorem 2 provides the exact probability computation for any number of nodes *p*. For instance, probability *P*(*D*_4_ ≥ 2.5) is computed iteratively as follows. We start with computing *A*_1,1_ = *A*_0,1_ + *A*_1,0_ = 2, *A*_2,1_ = *A*_1,1_ + *A*_1,0_ = 3, ⋯, *A*_4,4_ = *A*_4,3_ + *A*_3,4_ = 27 + 27 = 54 (red numbers in Fig. 2). Thus the probability is 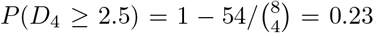. Few other examples that can be computed easily are

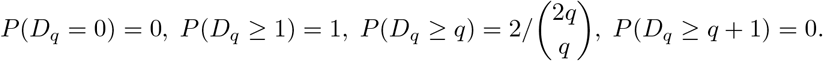

Computing *A_q,q_* iteratively requires at most *q*^2^ operations while permuting two samples consisting of q elements each requires 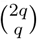 operations. Thus, our method can compute the *p*-value exactly substantially faster than the permutation test that is approximate and exponentially slow.

The asymptotic probability distribution of *D_q_* can be also determined for sufficiently large *q* without computing iteratively as Theorem 2.

### Theorem 3.

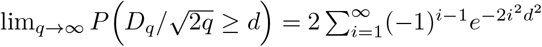.

The proof is not given here but the result follows from [8,14]. From Theorem 3, *p*-value under *H*_0_ is computed as

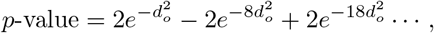

where *d_o_* is the *least integer greater than* 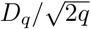 in the data (Fig. 3-left). For any large observed value *d*_0_ ≥ 2, the second term is in the order of 10^−14^ and insignificant. Even for small observed *d*_0_, the expansion converges quickly and 5 terms are sufficient.

**Fig. 3:**
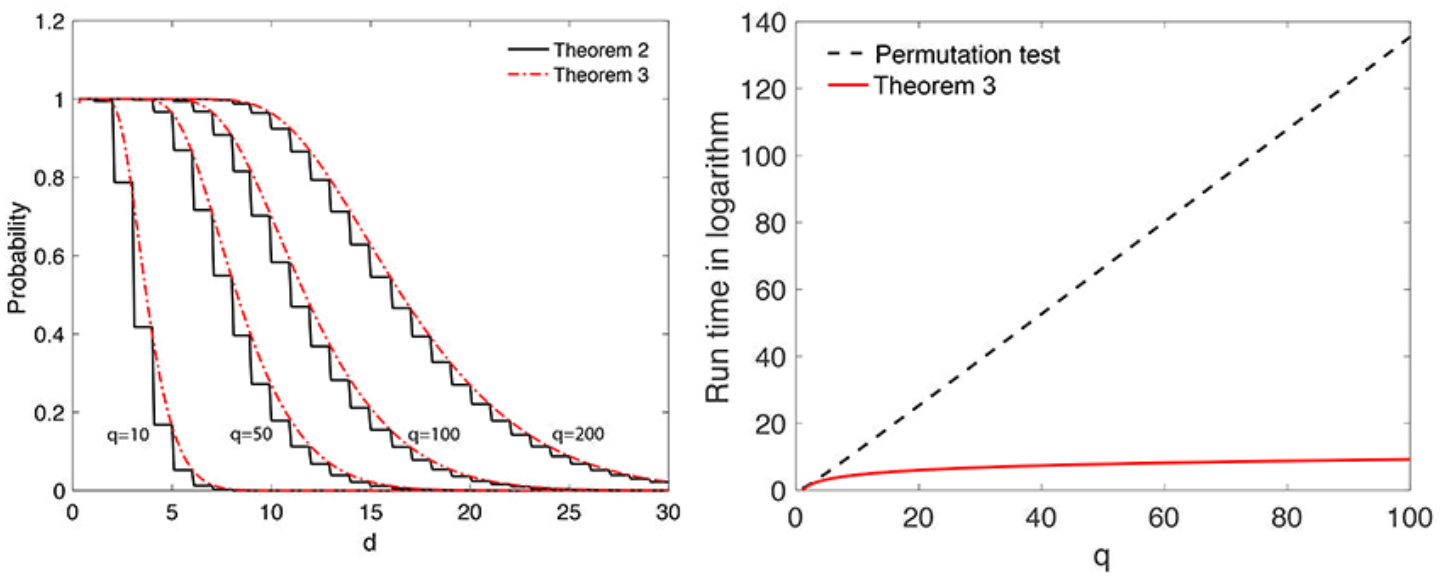
Left: Convergence of Theorem 2 to Theorem 3 for *q* = 10, 50, 100, 200. Right: Run time of permutation test (dotted black line) *vs*. the combinational method in Theorem 2 (solid red line) in logarithmic scale.

## 5 Validation and comparison

The proposed method was validated and compared in two simulation studies against Gromov-Hausdorff (GH) distance. GH-distance was originally used to measure distance between two metric spaces. It was later adapted to measure distances in persistent homology, dendrograms [3] and brain networks [11]. Following [11], the computation of GH-distance is done on graphs with1-correlation as edge weights. GH-distance is the difference in the *L*^∞^-norm of corresponding single linkage matrices.

### No network difference

The simulations were performed 1000 times and the average results were reported. There are three groups and the sample size is *n* = 5 in each group and the number of nodes are *p* = 40. In Groups I and II, data *x_k_*(*v_i_*) at each node *v_i_* was simulated as standard normal, i.e., *x_k_*(*v_i_*) ~ *N*(0, 1). The paired data was simulated as *y_k_*(*v_i_*) = *x_k_*(*v_i_*) + *N*(0, 0.02^2^) for all the nodes (Fig. 4). Using the proposed combinatorial method, we obtained the *p*-values of 0.6568 ± 0.3367 and 0.2830 ± 0.3636 for *β*_0_ and the size of the largest connected component demonstrating that the method did not detect network differences as expected.

**Fig. 4:**
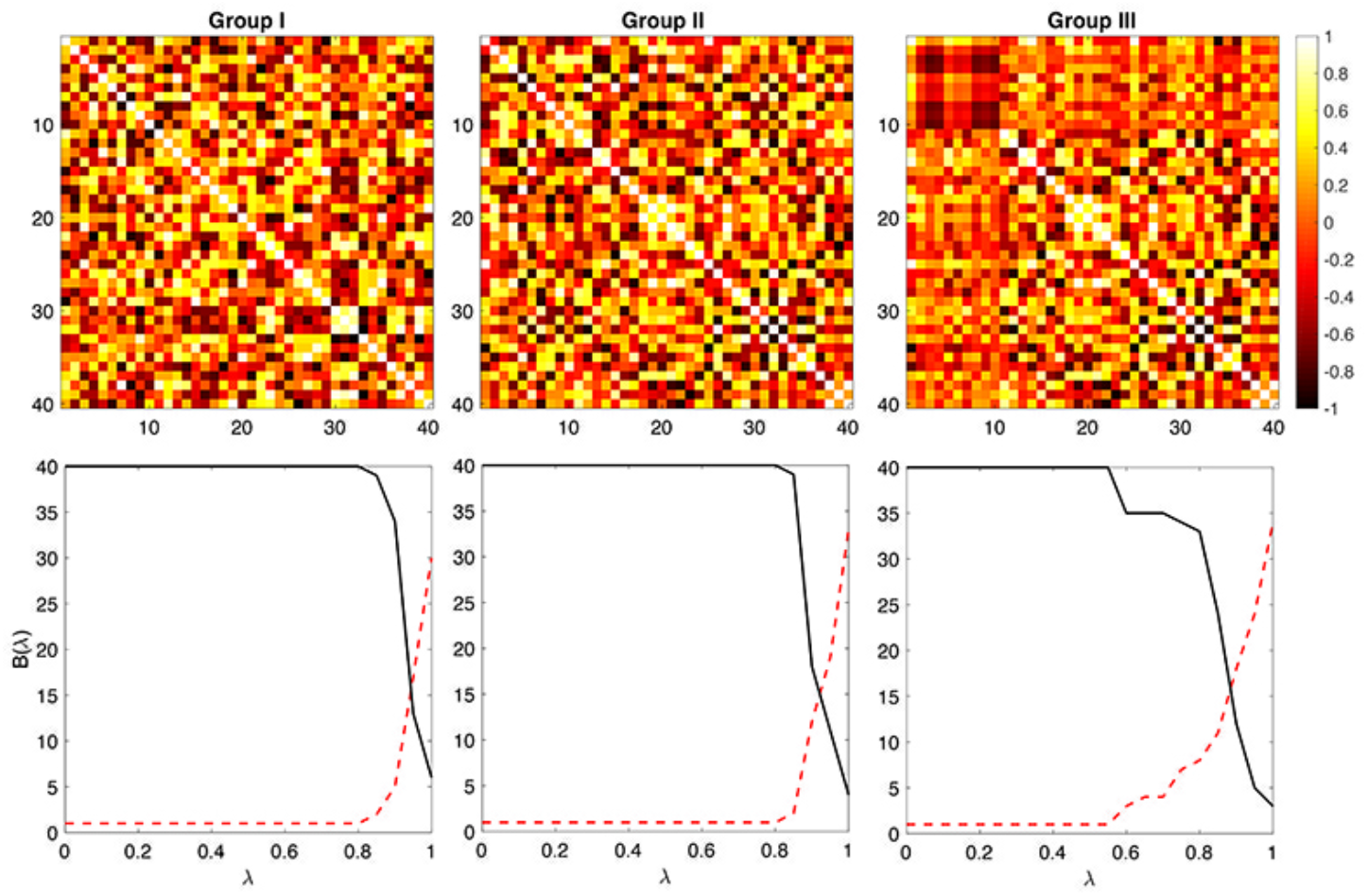
Simulation studies. Group I and Group II are generated independently and identically. The resulting *B*(λ) plots are similar. The dotted red line is *β*_0_ and the solid black line is the size of the largest connected component. No statistically significant differences can be found between Groups I and II. Group III is generated independently but identically as Group I but additional dependency is added for the first 10 nodes (square on the top left corner). The resulting B(λ) plots are statistically different between Groups I and II.

Using GH-distance, we obtained the *p*-value of 0.4938 ± 0.2919. The *p*-value was computed using the permutation test. Since there are 5 samples in each group, the total number of permutation is 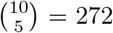 making the permutation test exact and the comparison fair. Both methods seem to perform reasonably well. However, GH-distance method took about 950 seconds (16 minutes) while the combinatorial method took about 20 seconds in a computer.

### Network difference

Group III was generated identically and independently like Group I but additional dependency was added by letting *y_k_*(*v_j_*) = *x_k_*(*v*_1_)/2 for ten nodes indexed by *j* = 1, 2, ⋯, 10. This dependency gives high connectivity differences between Groups I and III (Fig. 4). Using the proposed combinatorial method, we obtained the *p*-values of 0.0422 ± 0.0908 and 0.0375 ± 0.1056 for *β*_0_ and the size of the largest connected component. On the other hand, we obtained 0.1145 ± 0.1376 for GH-distance. The proposed method seems to perform better in the presence of signal. The `MATLAB` code for performing these simulations is given in http://www.cs.wisc.edu/~mchung/twins.

## 6 Twin fMRI study

The study consists of 11 monozygotic (MZ) and 9 same-sex dizygotic (DZ) twin pairs of 3T functional magnetic resonance images (fMRI) acquired in Intera-Achiava Phillips MRI scanners with a 32 channel SENSE head coil. Subjects completed monetary incentive delay task of 3 runs of 40 trials [10]. A total 120 trials consisting of 40 $0, 40 $1 and 40 $5 rewards were pseudo randomly split into 3 runs.

fMRI data went through spatial normalization to the template following the standard SPM pipeline. The fMRI dimensions after preprocessing are 53 × 63 × 46 and the voxel size is 3mm cube. There are a total of *p* = 25972 voxels in the template. After fitting a general linear model at each voxel, we obtained the contrast maps testing the significance of activity in the delay period for $5 trials relative to the control condition of $0 reward (Fig. 1). The paired contrast maps were then used as the initial images vectors **x** and **y** in the starting model (2).

We are interested in knowing *the extent of the genetic influence on the functional brain network* of these participants while anticipating the high reward as measured by activity during the delay that occurs between the reward cue and the target, and its statistical significance. The hyper-network approach is applied in extending the voxel-level univariate genetic feature called *heritability index* (HI) into a network-level multivariate feature. HI determines the amount of variation (in terms of percentage) due to genetic influence in a population. HI is often estimated using Falconer’s formula [6] as a baseline. MZ-twins share 100% of genes while DZ-twins share 50% of genes. Thus, the additive genetic factor A, the common environmental factor *C* for each twin type are related as

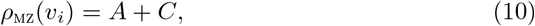

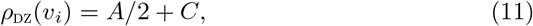

where *ρ_MZ_* and *ρ_DZ_* are the pairwise correlation between MZ- and and same-sex DZ-twins at voxel *v_i_*. Solving (10) and (11), we obtain the additive genetic factor, i.e., HI given by

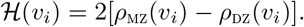

HI is a univariate feature measured at each voxel and does not quantify how the changes in one voxel is related to other voxels. We can determine HI across voxels *v_i_* and *v_j_* as

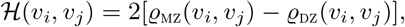

ϱ_MZ_ and ϱ_DZ_ are the symmetrized cross-correlations between voxels *v_i_* and *v_j_* for MZ- and DZ-twin pairs:

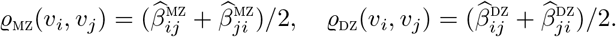

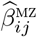 and 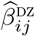 are the estimated cross-correlations from model (4). Note that

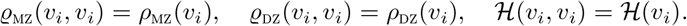

In Fig. 5-left and -middle, the nodes are *ρ*_MZ_(*v_i_*) and *ρ*_DZ_(*v_i_*) while the edges are ϱ_MZ_(*v_i_, v_j_*) and ϱ_DZ_(*v_i_, v_j_*) for λ = 0.5, 0.8. In Fig. 5-right, we have 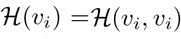. The network visualization shows high HI values for almost everywhere along the edges. We are interested in testing the statistical significance of the estimated HI for all edges. Testing for 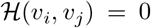 is equivalent to testing ϱ_MZ_(*v_i_, v_j_*) = ϱ_DZ_(*v_i_, v_j_*). Thus, we test for hyper-network differences between twins using the test statistic *D_q_* (*q* = 100 is used) (Fig. 6). For *β*_0_ and the size of the largest connected component *p*-values are less than 0.00002 and 0.00001 respectively indicating very strong significance of MZ- and DZ- network difference and HI of the whole brain network.

**Fig. 5:**
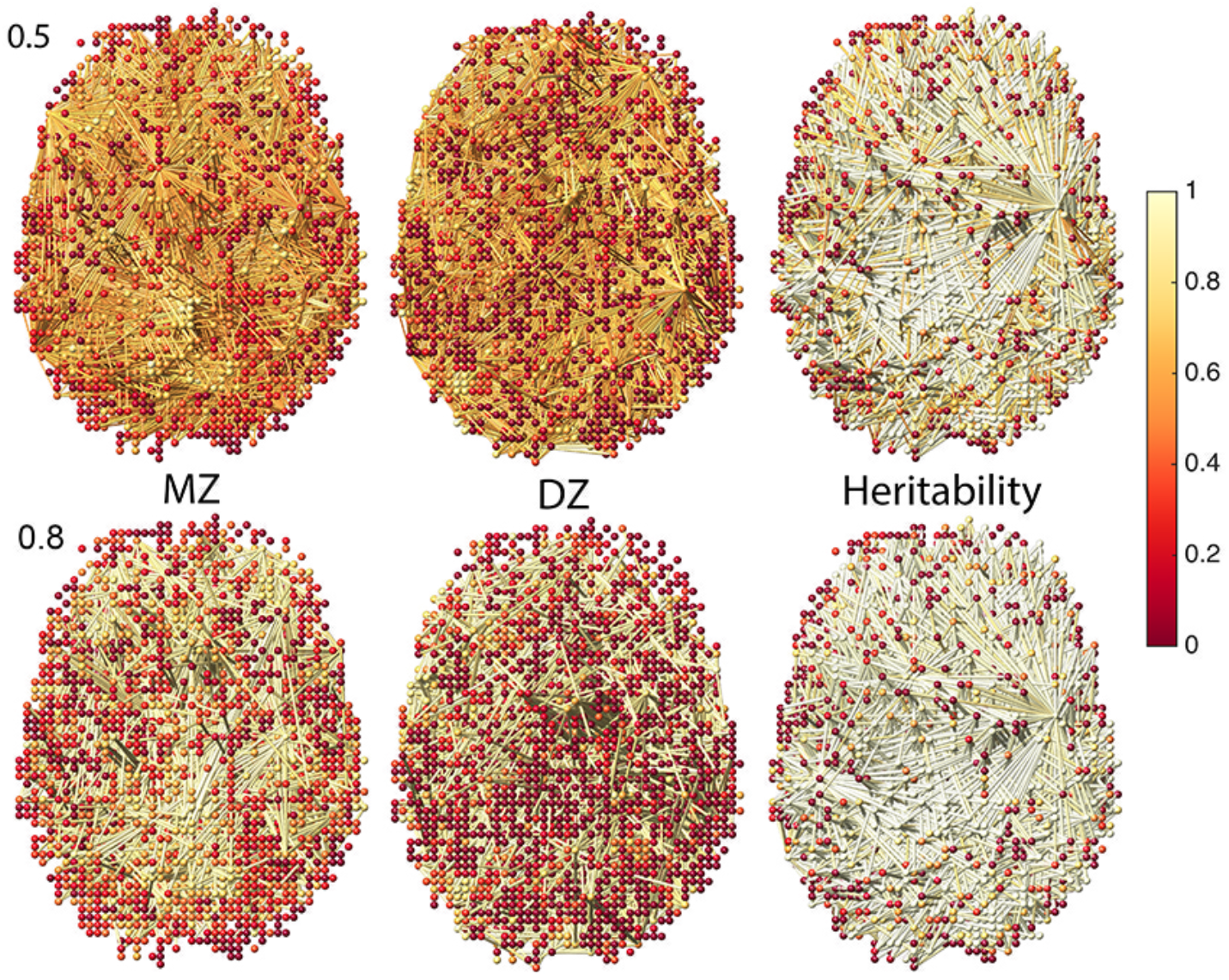
Left, middle: Node colors are correlations of MZ- and DZ-twins. Edge colors are sparse cross-correlations at sparsity λ = 0.5,0.8. Right: Heritability index (HI) at nodes and edges. MZ-twins show higher correlations compared to DZ-twins. Some low HI nodes show high HI edges. Using only the voxel-level HI feature, we may fail to detect such high-order genetic effects on the functional network.

**Fig. 6:**
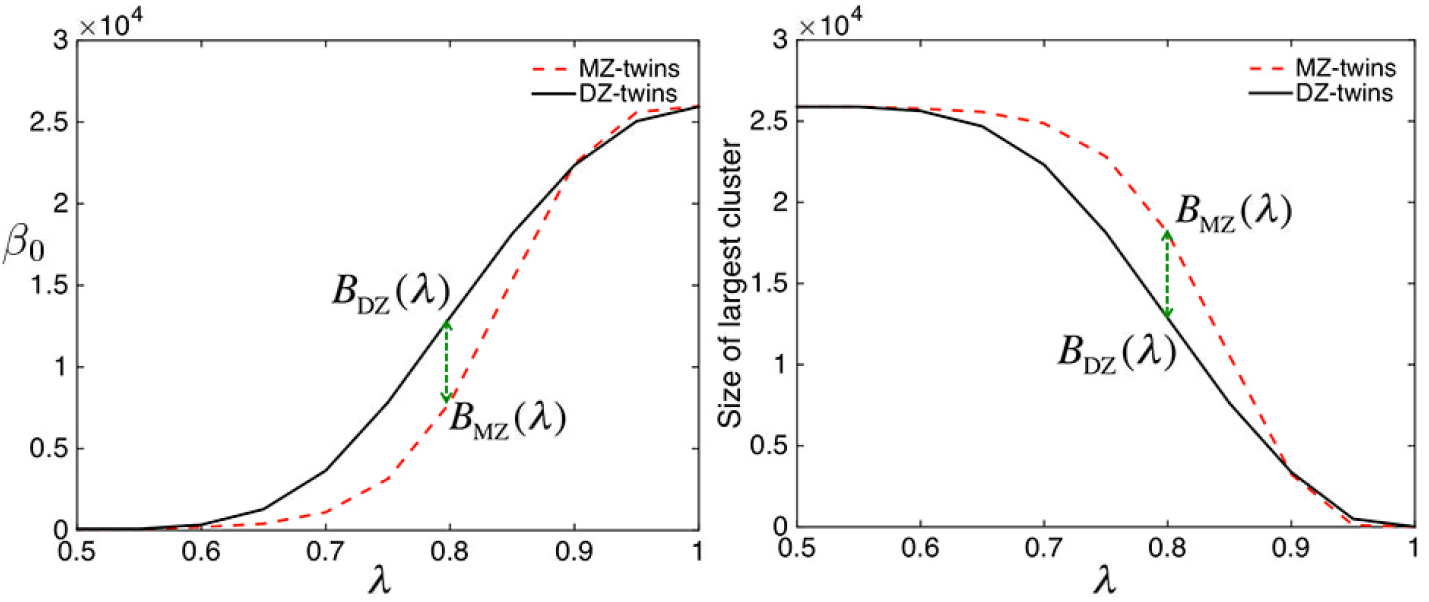
The result of graph filtrations on twin fMRI data. The number of connected components (left) and the size of the largest connected component (right) are plotted over the sparse parameter λ. For each λ, MZ-twins tend to have smaller number of connected components but larger connected component. The dotted green arrow (*D_q_*) where the maximum group separation occurs.

## 7 Discussion

### Hyper-network

In this paper, we presented a unified statistical framework for for analyzing paired images using a hyper-network. Although we applied the framework in twin fMRI, the method can be easily applied to other paired image settings such as longitudinal and multimodal studies. The method can be further extended to triple images. Then we should start with the following hyper-network model

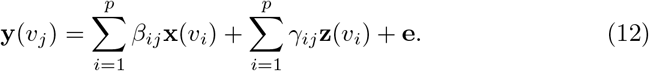

Since we usually have more number of nodes p than the number of images n, it is necessary to introduce two separate sparse parameters in (12). Then, instead of building graph filtration over 1D, we need to build it over 2D sparse parameter space. This is related to multidimensional persistent homology [4,12]. Extending the proposed method to triple image setting is left as a future study.

### Exact topological inference

We presented a combinatorial method for computing the probability that avoids time consuming permutation tests. Theorem 2 is the exact nonparametric procedure and does not assume any statistical distribution on graph features other than that they have to be monotonic. The technique is very general and applicable to any monotonic graph features such as node degree. Based on Stirling’s formula, 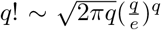, the total number of permutations in permuting two vectors of size *q* each is 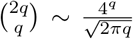. This is substantially larger than the quadratic run time *q*^2^ needed in our method (Fig. 3). Even for small *q* = 10, more than tens of thousands permutations are needed for the accurate estimation the *p*-value. On the other hand, only up to 100 iterations are needed in our combinatorial method.

## Acknowledgements

This work was supported by NIH grants 5 R01 MH098098 04 and EB022856 and NRF grant from Korea (NRF-2016R1D1A1B03935463). We thank Yoonsuck Choe of Texas A&M University and and Daniel Rowe of Marquette University for valuable discussions on permutation tests. We wish to thank anonymous reviewers for valuable comments that improved the revision.

